# Complete Workflow for High Throughput Human Single Skeletal Muscle Fiber Proteomics

**DOI:** 10.1101/2023.02.23.529600

**Authors:** Amanda Momenzadeh, Yuming Jiang, Simion Kreimer, Laura E. Teigen, Carlos S. Zepeda, Ali Haghani, Mitra Mastali, Yang Song, Alexandre Hutton, Sarah J Parker, Jennifer E. Van Eyk, Christopher W. Sundberg, Jesse G. Meyer

## Abstract

Skeletal muscle is a major regulatory tissue of whole-body metabolism and is composed of a diverse mixture of cell (fiber) types. Aging and several diseases differentially affect the various fiber types, and therefore, investigating the changes in the proteome in a fiber-type specific manner is essential. Recent breakthroughs in isolated single muscle fiber proteomics have started to reveal heterogeneity among fibers. However, existing procedures are slow and laborious requiring two hours of mass spectrometry time per single muscle fiber; 50 fibers would take approximately four days to analyze. Thus, to capture the high variability in fibers both within and between individuals requires advancements in high throughput single muscle fiber proteomics. Here we use a single cell proteomics method to enable quantification of single muscle fiber proteomes in 15 minutes total instrument time. As proof of concept, we present data from 53 isolated skeletal muscle fibers obtained from two healthy individuals analyzed in 13.25 hours. Adapting single cell data analysis techniques to integrate the data, we can reliably separate type 1 and 2A fibers. Sixty-five proteins were statistically different between clusters indicating alteration of proteins involved in fatty acid oxidation, muscle structure and regulation. Our results indicate that this method is significantly faster than prior single fiber methods in both data collection and sample preparation while maintaining sufficient proteome depth. We anticipate this assay will enable future studies of single muscle fibers across hundreds of individuals, which has not been possible previously due to limitations in throughput.

## INTRODUCTION

Skeletal muscle (skm), which comprises ∼40% of human body mass, is characterized by remarkable heterogeneity of cell types that allows muscles to perform a wide variety of functional tasks. Three major skm cell types – commonly known as fibers – exist in human limb muscles that are differentiated primarily by their contractile kinetics from slow (type 1) to fast (type 2A and 2X) and metabolic profiles from more oxidative phosphorylation to more glycolytic ^1,2^. Slow fibers are involved primarily with low-intensity, continuous contractions, such as the maintenance of posture or walking, while fast fibers allow for rapid, forceful contraction. The differences in contractile kinetics between the fiber types are determined in large part by the myosin heavy chain (MYH) isoform that is expressed, namely MYH7 (type 1), MYH2 (type 2A), and MYH1 (type 2X)^3^. However, the fiber type diversity in human skm expands well beyond the contractile kinetics from the different MYH isoforms to include proteins involved in cell structure, excitation contraction coupling, regulatory proteins, and metabolism^1^. Thus, to capture the cellular diversity in human skm and elucidate how the muscle adapts to acute and chronic perturbations (e.g., exercise training, bedrest, disease, or aging) in a fiber-type specific manner requires technological advancements in large-scale single cell proteomics analysis.

Recent breakthroughs in single fiber proteomics have allowed for isolation of proteins found solely within the skm fiber, without the contamination from adipocytes, connective tissue, blood vessels and nerve cells that are all present in human skm biopsy lysates^3^. These studies have overcome many of the major factors that complicate proteomic analysis of skm fibers, such as the predominance of the fiber protein mass taken up by the contractile proteins (myosin and actin) and the large variability in the size of fibers both within and across individuals ^3–5^, allowing for several key discoveries to be made in human skm biology. For example, it was found that the aging-related decrements in the size and contractile function of the fast fibers in older adults^6^ may be explained, in part, by a downregulation of proteins involved in sarcomere homeostasis and protein quality control of this fiber type^7^. However, despite these recent successes, the current methods used in single fiber proteomics analysis are slow and laborious, limiting the number of fibers that can be studied per individual and making population scale single fiber proteomics studies impossible. Most previous studies also used MaxQuant’s match-between-runs (MBR) feature from larger muscle biopsy lysate analyses, which addresses missing peptides by allowing peptide peaks to be transferred between runs^8^. However, without false identification transfer control, MBR may introduce false positives at a high rate, compromising protein quantification accuracy^8,9^.

Here we developed a method for high throughput single muscle fiber proteomics that we applied to 53 human single skm fibers obtained from the vastus lateralis of two female volunteers. Compared to prior single muscle fiber proteomics publications, our method uses streamlined proteomic sample preparation that does not require desalting along with MS data collection at a rate of 96 samples per day. Additionally, previous single skm proteomics papers have used Oribitrap mass spectrometers (ThermoScientific)^3,5,7,10^. In this study, we used a timsTOF SCP (Bruker) to collect diaPASEF^11^ data from single skm fibers, allowing nearly a 10 fold increase in throughout while still quantifying roughly the same number of proteins per single fiber compared to when MBR in MaxQuant was not used (see supplementary figure 2 in Murgia, et al. 2017)^7^. Additionally, we apply the python data analysis package (scanpy), a tool designed for single-cell transcriptomics data, to perform unsupervised clustering of single skm fiber proteomics data^11,12^. Our new workflow including sample preparation, data collection, and bioinformatics provides a rapid and complete platform for single muscle fiber proteomics that will enable large cohort studies of muscle proteome composition in the future.

## METHODS

### Participants

Two female volunteers (22 and 69 years) participated in this study. Participants were self-reported to be healthy, community dwelling adults free of any known neurological, musculoskeletal, or cardiovascular diseases. Participants provided written informed consent, and procedures were approved by the Marquette University Institutional Review Board and conformed to the principles in the Declaration of Helsinki.

### Skeletal Muscle Biopsy

Percutaneous muscle biopsies were obtained from the vastus lateralis using the Bergstrom needle technique^14^. Participants were instructed to abstain from strenuous exercise for 48 h prior to the biopsy procedure and to arrive at the laboratory fasted and without the consumption of caffeine. The biopsy site was cleaned with 70% ethanol, sterilized with 10% povidone-iodine and anesthetized with 1% lidocaine HCl. A small ∼1 cm incision was made overlying the distal one-third of the muscle belly, and the biopsy needle was inserted under local suction to obtain the tissue sample as described previously^15^. Fiber bundles from the biopsy were separated, arranged longitudinally on a small notecard, frozen in liquid nitrogen-cooled isopentane (AC126470250; Acros Organics) and subsequently stored at -80°C.

### Single Muscle Fiber Isolation

Skeletal muscle tissue samples were allowed to thaw in a standard petri dish coated with Sylgard containing a cold (∼4°C) relaxing solution commonly used in single fiber contractile function experiments^15,16^ but with the addition of a protease and phosphatase inhibitor cocktail (78440; Thermo Scientific). The composition of the solution was 79.2 mM KCl, 20 mM imidazole, 7 mM EGTA, 5.44 mM MgCl_2_, 4.735 mM Na_2_ATP, and 14.5 mM Na_2_ phosphocreatine, and the pH of the solution adjusted to 7.0 with KOH. Single muscle fibers were manually isolated under a dissecting microscope with fine tipped tweezers. Individual fibers were then carefully transferred into separate wells of a 384-well PCR plate (HSP3805; BioRad). Both the petri dish and the 384-well PCR plate were maintained at ∼4°C during the muscle fiber isolation process with cold plates. The 384-well PCR plate was covered with aluminum sealing foil (MSF1001; BioRad) and stored at -80°C until further analysis.

### Sample Preparation

Four hundred nanoliters lysis buffer containing 100mM triethylammonium bicarbonate (TEAB; 18597, Sigma; pH 8.5), 0.2% DDM, 10ng/ul Trypsin (Promega) was dispensed into each well using the cellenONE system (cellenONE®, Cellenion). After freezing and thawing for 5 minutes in -80°C, 400nl of trypsin 50 ng/ul in 60mM TEAB was dispensed into each well again and the plate was incubated at 37°C for 3.5 hours. Digestion was quenched with 100nL 1% aqueous formic acid. The plate was stored at -80°C until MS analysis.

### Liquid Chromatography

Dual trap single column (DTSC)^17^ was adapted for nanoflow to enable single cell analysis^18^. The trapping columns were 0.17 µL media bed (EXP2 from Optimize Technologies) packed with 10 µm diameter, 100 Å pore PLRP-S (Agilent) beads, and the analytical column was a PepSep 15 cm × 75 µm packed with 1.9 µm C18 stationary phase (Bruker). The configuration was installed on a Thermo Ultimate 3000 nanoRSLC equipped with one 10-port valve and one 6-port valve and a nano flow selector. Mobile phase A as 0.1% formic acid in water. Mobile phase B was 0.1% formic acid in acetonitrile. Peptides were separated using the following gradient all using linear transitions between conditions: starting conditions of 9% B at 500 nL/min; 22% B over 8 minutes; 37% B over 4.7 minutes; 1000 nL/min flowrate and 98% B over 0.2 minutes; hold at 98% B for 1 minute; reduce to 9% B at 1,000 nL/min over 0.1 minutes; hold at 9% B at 1,000 nL/min for 0.9 min; return to 500 nL/min flowrate in 0.1 min (15 min total). Valves and trapping columns were heated to 55° C and the analytical column was heated to 60° C. See Kreimer et al. for additional details about the dual trap chromatography setup.^18^

### Mass Spectrometry

The analytical column was directly connected to a 10 µm ZDV emitter (Bruker) inside of the Bruker captive source. The capillary voltage was set to 1700 V with the dry gas set at 3.0 L/min and 200° C. Data was acquired by diaPASEF on a Bruker timsTOF SCP with the ion accumulation and trapped ion mobility ramp set to 166 ms. DIA scans were optimized and acquired with 90 *m/z* windows spanning 300 to 1200 m/z and 0.6 to 1.43 1/K0. A 0.86 s cycle time included one full MS1 scan followed by 4 trapped ion mobility ramps to fragment all ions within the defined region.

### Mass Spectrometry Data Analysis

#### Peptide and protein quantification

A spectral library was generated by a library-free search in DIA-NN 1.8^19,20^ using a FASTA of the human proteome with one entry per gene downloaded on January 5, 2023 with 20,594 entries. The library from that first library-free search was used to analyze MS data from 53 single skm fibers each containing more than 500 proteins per fiber in the first search. Protein identifications by DIA-NN are based on unique peptides. The option for re-annotation of identified peptides with the FASTA was used. The targeted library from the second search contained 1,796 protein groups and 11,610 precursors in 9,790 elution groups. Protein groups are formed when identified peptides cannot differentiate between multiple proteins with shared subsequences. Each search was conducted with C carbamidomethylation off, match-between-runs (MBR) enabled, double pass mode enabled, cross-run normalization disabled, speed and RAM usage set to optimal, and any LC (high-accuracy) quantification strategy. MBR in DIA-NN is not equivalent to MBR in MaxQuant; in DIA-NN, MBR means the spectral library for the first search is used for the second search. Precursor charge range was set between 2 and 4, mass accuracy fixed to 15 ppm, and one missed cleavage was allowed in the database search. All other parameters were the default. Intensity based absolute quantification (iBAQ)^21^ was manually implemented in python and used to calculate protein group quantities from peptide quantities. Peak intensities for all peptides corresponding to a protein group were summed and then divided by the number of possible unique peptides, which was calculated to allow for peptides with one missed cleavage and between 7 to 30 amino acids long to mirror DIA-NN settings. Resulting protein quantities in each well were normalized to alpha skeletal muscle actin (ACTS, P68133) quantity in that well to control for fiber length and cross-sectional area, as previously described^3,5^. Lastly, fibers were ranked by percent of their dominant MYH isoform. Pure MYH7, MYH2 or MYH1 myofibers were assigned if any of these three isoforms reached 80% of the total MYH signal for all isoforms^3^. Fibers that did not have a single MYH at more than 80% of one specific isoform were labeled as one of two hybrids based on the top two most prevalent MYHs, either co-expression of MYH1 and MYH2 or MYH2 and MYH7. Protein intensities from MaxLFQ^22^ were also normalized to the quantity of ACTS in each well. An example extracted ion chromatogram for a random peptide was obtained from Skyline-daily^23^ version 22.2.1.351 to show the quality of chromatography. All data analysis was performed in python version 3.9.12.

#### Data Visualization

Using scanpy version 1.9.1^24^, we implemented a pipeline using Uniform Manifold Approximation & Projection (UMAP)^25^ and the Leiden method^26^ to reduce the data to a two-dimensional space and bin the data into discrete populations, respectively. Proteins appearing in less than 5% of fibers (or a minimum of 3 fibers) were filtered out as these are rare. To obtain parameters leading to the best match to manually annotated cluster labels, we used a random search with 300 iterations optimizing number of principal components (PCs) that explain variance (range 1-50), neighbors that compute a cell’s local connectivity (range 1-15 or 30), Leiden resolution which controls coarseness of clustering (range 0-1 by a factor), and whether to scale data to unit variance and zero mean. Using a variety of metrics, we measured similarity between the Leiden inferred cluster labels and author annotated myosin labels based on MYH percentage found in fibers associated with each cluster. The set of parameters with the highest normalized mutual information score (NMI) was used for the final UMAP and Leiden clustering. NMI, which ranges from 0 (similarity expected by chance) to 1 (perfect similarity), provides a measure of quality of clustering and allows for comparison between datasets with different numbers of clusters. Multiple sets of parameters led to the same optimized NMI score of 0.7, and one set was chosen at random (no scaling, 33 PCs, 11 neighbors, 0.1 resolution). We also reported conditional entropy, which refers to the amount of information needed to describe a reference label given the Leiden clustering generated labels (low value means Leiden labels are very informative of reference labels) corresponding to the parameters with the highest NMI. A Chi squared test was used to test for a difference between young and old fibers between clusters.

#### Pathway Enrichment Analysis

We first determined proteins that were most uniquely identifying clusters. Pairwise protein distribution comparisons were made between Leiden groups using a Wilcoxon Rank-Sum test and Benjamini-Hochberg (B-H) multiple testing correction with alpha=0.01. Gene ontology (GO) Biological Process term enrichment analysis was applied to proteins with an adjusted p-value less than 0.01 (negative log_10_ adjusted p-value >2) and an absolute value of log_2_ fold change of at least 1 between Leiden groups. Term enrichment analysis was performed using ClueGO (version 2.5.9)^27^ application within Cytoscape (version 3.9.1; release date 5/25/2022)^28^. WikiPathways term enrichment was used to understand the most common pathways among all 744 proteins and GO Biological Process using GO term fusion was used for network analysis of differentially expressed proteins between Leiden clusters. Only pathways with a p-value <0.0001 were enriched for either pathway. For either analysis, we manually removed redundant terms to keep only those that connected all the proteins to the simplest network.

### Code and Data Availability

All mass spectrometer raw data, libraries and outputs have been added to MassIVE database: doi:10.25345/C54T6FC9Z (password: hello). JupyterLab notebooks can be found at xomicsdatascience/HT-Skm-Proteomics (github.com).

## RESULTS

Our high throughput single skm fiber proteomics workflow enabled collection of high quality and high content protein data from 53 human fibers (31 from young and 22 from old), obtained from the vastus lateralis of two female volunteers (**Figure 1A**). Single fibers were dissected into wells of 384 well plates, and then one pot sample preparation was carried out before directly loading 1/100th of the total sample from the same plate onto a timsTOF SCP MS for a 15-minute total run time with diaPASEF data collection.

**Figure 1.**
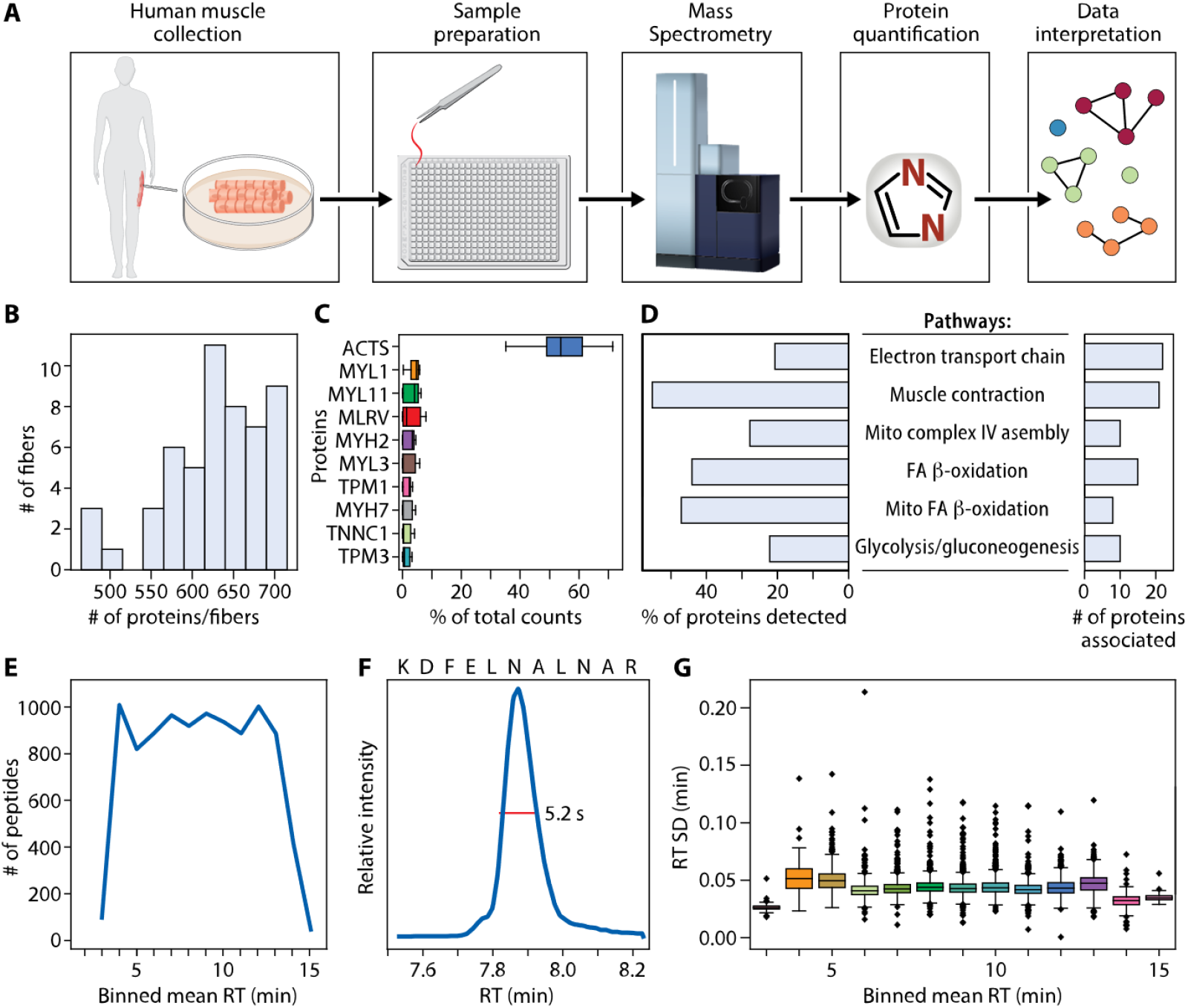
Data and workflow overview. **(A)** Muscle biopsies were collected from two healthy female donors. Fibers were processed using one pot in-solution digestion in a 384 well plate and data was acquired by DIA-PASEF on a Bruker timsTOF SCP. Proteins were quantified using library-free search with DIA-NN against the whole human proteome. Clustering allowed for visualization of similarly grouped fibers. Top ranked proteins distinguishing clusters were then used to understand biological pathways. **(B)** Number of proteins quantified per fiber. **(C)** Ten proteins with highest percent of total counts in each fiber. **(D)** WikiPathways term enrichment analysis summary of all 744 proteins identified in any fiber. (**E)** Number peptides quantified per minute across binned mean retention times. **(F)** Extracted ion chromatogram showing a typical full width at half maximum (FWHM) of 5.2 seconds for example peptide KDFELNALNAR. **(G)** RT standard deviation in minutes across binned mean RTs.

At a false discovery rate (FDR) of 1%, we identified 868 unique protein groups present in at least one fiber. There were 124 protein groups with multiple members, which were filtered out to simplify subsequent analyses, resulting in 744 unique proteins (**Table S1**) corresponding to 8,764 unique precursors. Proteins were included if they were identified in at least 3 fibers (>5%). An average of 629 ± 62 proteins were quantified per single muscle fiber (**Figure 1B**). Forty-one percent (305/744) of proteins were quantified in all 53 fibers. The most abundant protein across all fibers was ACTS (**Figure 1C**). WikiPathways enriched in the set of all proteins is depicted in **Figure 1D**; proteins involved with striated muscle contraction composed the largest group of proteins (55%) detected from a pathway compared to all the proteins associated with that pathway, followed by mitochondrial fatty-acid beta-oxidation (47%). Our chromatography method showed consistent instrument utilization across the elution gradient with more than 800 peptides quantified in each minute of data collection between 4 and 13 minutes (**Figure 1E**). Our chromatographic peaks were sharp with an example extracted ion chromatogram for a random peptide sequence (KDFELNALNAR) showing a full width at half maximum (FWHM) of 5.2 seconds. Retention time (RT) standard deviations were small across binned mean RTs (**Figure 1G**).

The 10 most abundant iBAQ-calculated proteins in pure type 2A or 1 fibers are plotted separately **(Figure 2A-B)**, demonstrating the high abundance of ACTS and congruence of regulatory protein isoforms with their appropriate primary MYH isoform. Although MYL1 (a fast myosin regulatory light chain) was present among the top 10 most abundant proteins in both fiber types, its quantity was two-fold higher in type 2A fibers (**Figure 2C**). Among the proteins with top 10 highest percentages of total counts per fiber from MaxLFQ quantification, ACTS was not as clearly more abundant (**Figure S1**).

**Figure 2.**
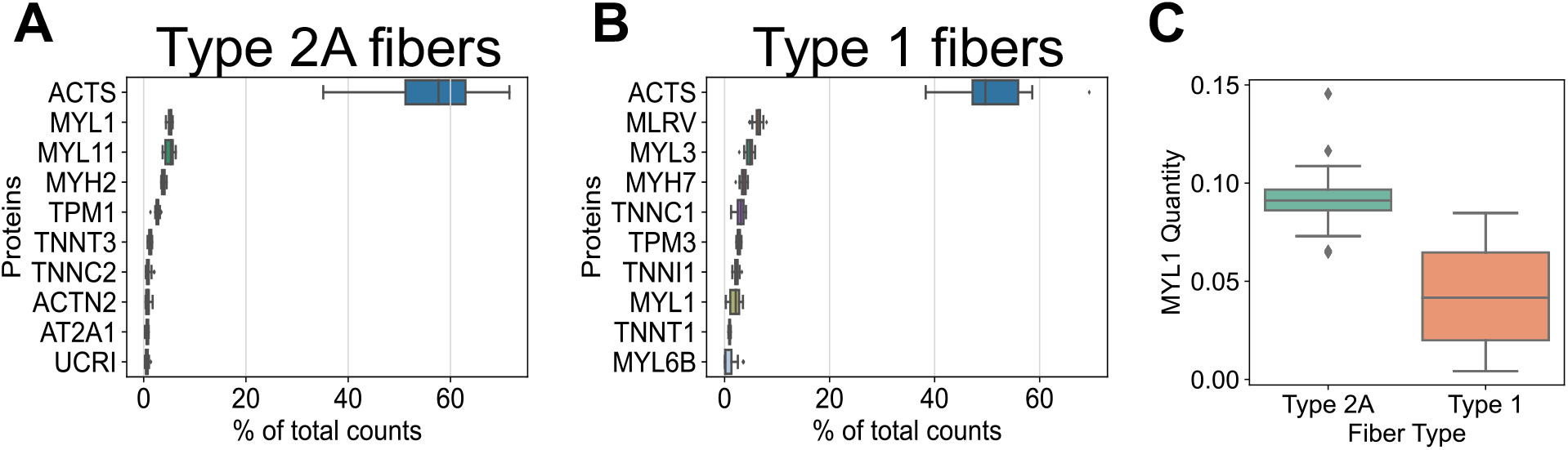
Ten most abundant proteins when quantities are computed by iBAQ. **(A)** Top ten proteins with highest percent of total counts in pure type 2A fibers quantified using iBAQ. **(B)** Top ten proteins with highest percent of total counts in pure type 1 fibers quantified using iBAQ. **(C)** MYL1 iBAQ-calculated quantity in type 1 versus 2A fibers. MLRV is interchangeable with MYL2. MYL11 is interchangeable with MYLPF and MLC2-fast

We explored differences in ratios of MYH proteins per fiber between iBAQ and MaxLFQ, which are different protein quantification algorithms. A fiber was assigned as a pure type 1 or 2A if the fraction of MYH7 or MYH2, respectively, of all MYH isoforms combined was greater than 80%. Fibers not meeting this threshold were categorized as mixed fibers based on the top two most prevalent MYHs. Using MaxLFQ and a 80% cutoff for identifying pure fibers, there were 17 MYH7, 3 MYH1, and only 1 MYH2 fiber. The mixed fibers consisted of 5 MYH2/7, 6 MYH1/2, 1 MYH1/7 and the remaining 20 fibers had a combination of MYH2 and MYH4 (**Figure 3A, Table S2**). The number of fibers expressing MYH4 is shown in **Figure 3B**. Initially there was one peptide (DEELDQLKR^2+^) mapped to MYH4 that differed from MYH2 by the assignment of L versus I, so this peptide was removed before subsequent iBAQ calculations (**Figure S2**). Using DIA-NN with iBAQ calculated protein intensities, we found that the majority of fibers had a single predominant MYH isoform (26 MYH2 and 18 MYH7) and the rest were mixed type (5 MYH2/7 and 4 MYH1/2) (**Figure 3C, Table S3**). Expression of MYH4 (**Figure 3D**), which is thought to be found in rodent skm and only in a specialized muscle type in humans^29^, was 0.3% of total MYH content when protein quantities were computed using iBAQ, but 10% of total MYH when using MaxLFQ derived protein quantities. Other single cell human skm studies also observe negligible or no MYH4 present in fibers^30^. MYH6, which is found in cardiac muscle cells^31^, was low in both quantification strategies–0.3% of total MYH content when protein quantities were calculated using iBAQ and 0.8% of total MYH content when using MaxLFQ. The drastic reduction in MYH2 fibers, the unlikely MYH1/7 hybrid, and a large number of fibers expressing MYH4 (**Figure 3B**) indicate that protein quantities computed with iBAQ are preferred over those from MaxLFQ when computing ratios of proteins.

**Figure 3.**
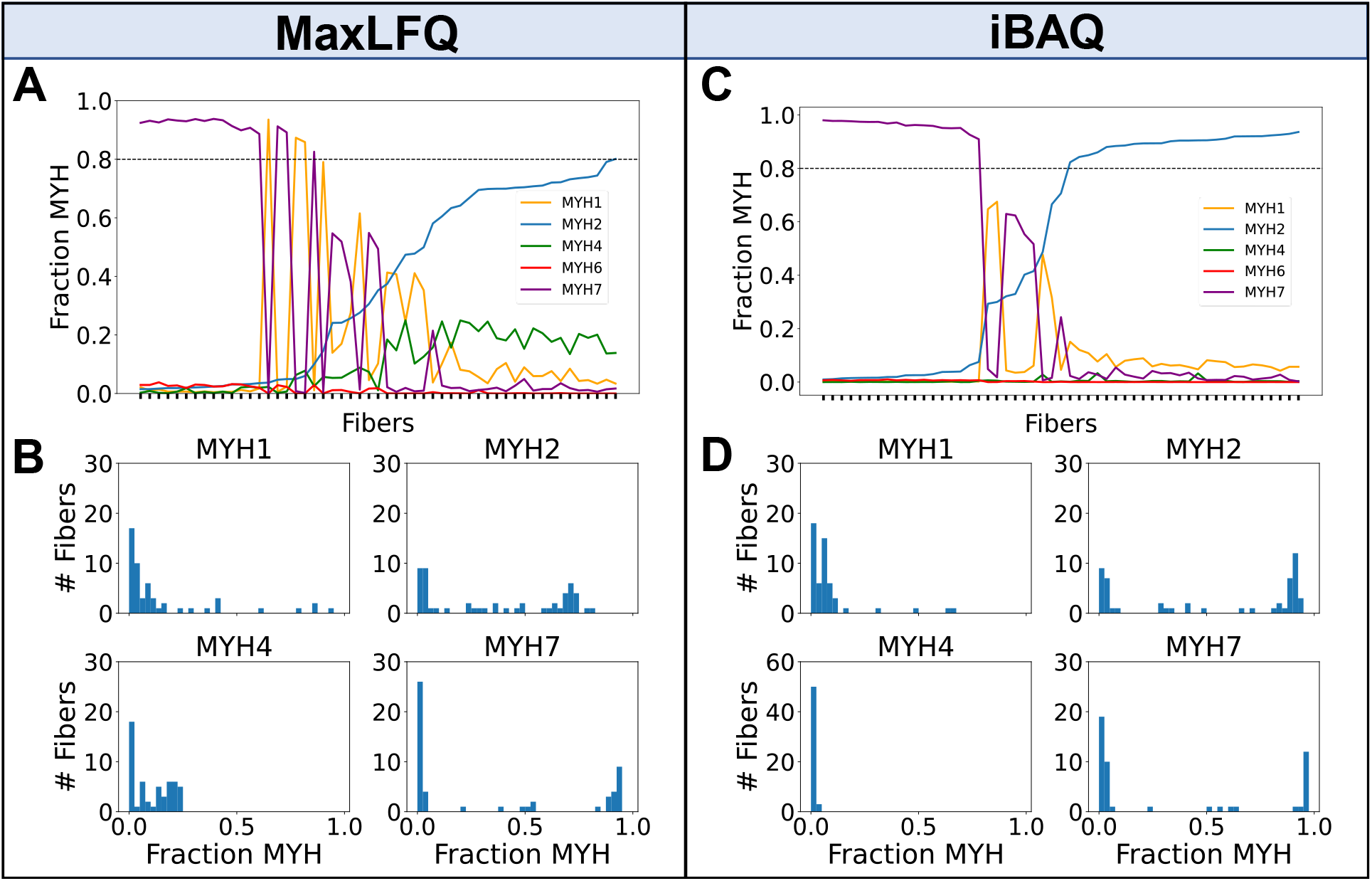
Detected proportions of MYH isoforms across 53 skeletal muscle fibers. **(A)** Fraction of MYH proteins per fiber (sorted by lowest to highest MYH2 quantity from left to right across the 53 fibers) derived from MaxLFQ. A horizontal line drawn at MYH fraction of 0.8 indicates the cut-off for assigning pure fibers. **(B)** Histogram of fibers per MYH fraction for each subtype using MaxLFQ. **(C)** Fraction of MYH proteins per fiber using protein quantities calculated using iBAQ. **(D)** Histogram of fibers per MYH fraction for each subtype using protein quantities calculated using iBAQ. MYH4 y-axis limits are different from rest of panels due to much greater proportion of zeros.

We performed unsupervised Leiden clustering^26^ and visualization with UMAP using all proteins as input to better understand how proteome profiles would group the fibers. Given the manual groupings defined by MYH proportions, we optimized the data reduction and clustering to best match those groups. Three hundred random combinations of workflow parameters were tested with a random search (see Methods). We used NMI to measure the similarity between the data inferred labels and the author assigned labels. The parameters resulting in the highest NMI of 0.7 (no data scaling, 33 PCs, 11 neighbors, 0.1 resolution) were used for the final clustering and UMAP plot. The corresponding conditional entropy was 0.72. We did not expect perfect similarity scores because all proteins were used to generate the UMAP plots and the manual annotations are only based on four MYH subtypes.

The optimal Leiden clustering separated all protein level data into two clusters (**Figure 4A**). When colored by the manual MYH annotations, we see clear separation between pure MYH2 and MYH7 fibers, as well as mixed MYH2/7 and MYH1/2 fibers (**Figure 4B**). When comparing the two UMAP plots, Leiden cluster 1 aligns closely with MYH7 and MYH2/7 annotated fibers, and Leiden cluster 0 with both MYH2 and MYH1/2 annotated fibers. **Figure 4C** shows protein identifications per fiber are random within clusters (i.e., a cluster is not driven by a high or low count). The UMAP plot of fibers colored by iBAQ-calculated abundance of MYH1, 2 and 7 (**Figure 4D**) aligns with the manual MYH annotated fiber groups in **Figure 3B**.

**Figure 4.**
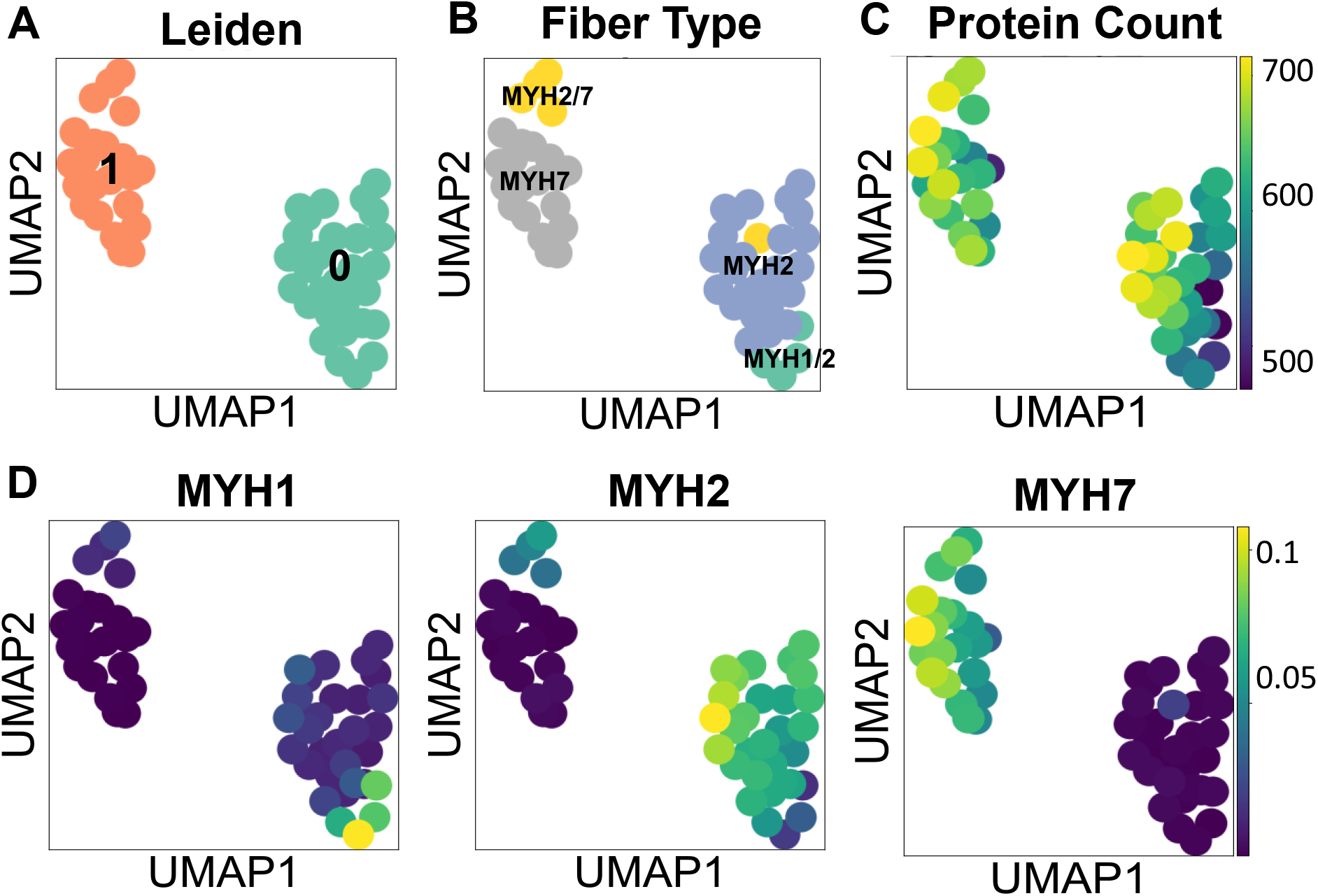
Clustering and dimension reduction of single fiber proteome data. **(A)** UMAP colored by Leiden clusters. **(B)** UMAP colored by author annotated fiber labels. **(C)** UMAP colored by the number of proteins identified per fiber. **(D)** UMAP colored by iBAQ-calculated quantities of MYH1, MYH2, MYH7 proteins.

Next, we performed a B-H corrected Wilcoxon Rank-Sum test with p-value <0.01 and a log_2_ fold change of at least 1 as the cutoff to identify differentially expressed proteins between Leiden clusters 0 and 1 (**Figure S4**). Sixty-five proteins were differentially expressed, of which 16 were upregulated in Leiden cluster 1 (mostly MYH7 fibers) and 49 were higher in Leiden cluster 0 (mostly MYH2 fibers). There was no difference between young and old fiber distribution between Leiden clusters (65% young in Leiden 0 and 50% young in Leiden 1; p-value=0.4), and although we are underpowered to compare fiber ratios between people, we note that four out of the five MYH2/7 hybrid fibers came from the older volunteer (**Figure S5**).

GO biological pathway enrichment using the 65 differentially expressed proteins (**Table S4**) between Leiden 0 and 1 clusters revealed proteins associated with two main pathways, muscle contraction and fatty acid beta-oxidation (**Figure 5A**). Many of these were canonical differences between fiber types, including skeletal troponin subunits isoforms (TNNI1, TNNT1, TNNC1 higher in slow fibers and TNNC2, TNNI2, TNNT3 higher in fast fibers), tropomyosins (TPM3 higher in slow fibers and TPM1 higher in fast fibers), myosin light chain proteins (MYL3, MYL5, MYL6B higher in slow fibers and MYL11, also known as MYLPF or MLC2-fast, higher in fast fibers), Z-line linking proteins (MYOZ2 higher in slow fibers and ACTN3 higher in fast fibers) and a sarcoplasmic reticulum protein (CASQ2 higher in slow fibers). Tropomyosins TPM1 and TPM3, which regulate muscle contraction, were both present in every fiber despite their expected specificity for fast and slow fibers, respectively. Analysis of their relative proportions showed two groups where one isoform was dominant over the other (**Figure S3A**). Correlation between the fast and slow myosin and tropomyosin isoforms was assessed. The fast form of myosin (MYH2) was well correlated with the fast form of tropomyosin (TPM1) while the slow myosin (MYH7) was well correlated with the slow form of tropomyosin (TPM3). There was minimal correlation when comparing the quantities of mismatched fast and slow myosins and tropomyosins (**Figure S3B**), overall supporting the quality of protein quantification with our method. All proteins associated with fatty acid beta-oxidation had higher expression in the Leiden 1 cluster. Example quantity distributions of log_2_ transformed iBAQ-calculated quantities for three proteins from fatty acid beta-oxidation pathway, MCAT, ACAT1, and ACADS, are visualized using raincloud plots (**Figure 5B**). Although this term enrichment analysis indicates MYH14 (a non-muscle MYH) was different between groups, MYH14 comprised 0.008% of total MYH content and the highest MYH14 quantity in any fiber was 0.02% of total MYH in that fiber.

**Figure 5.**
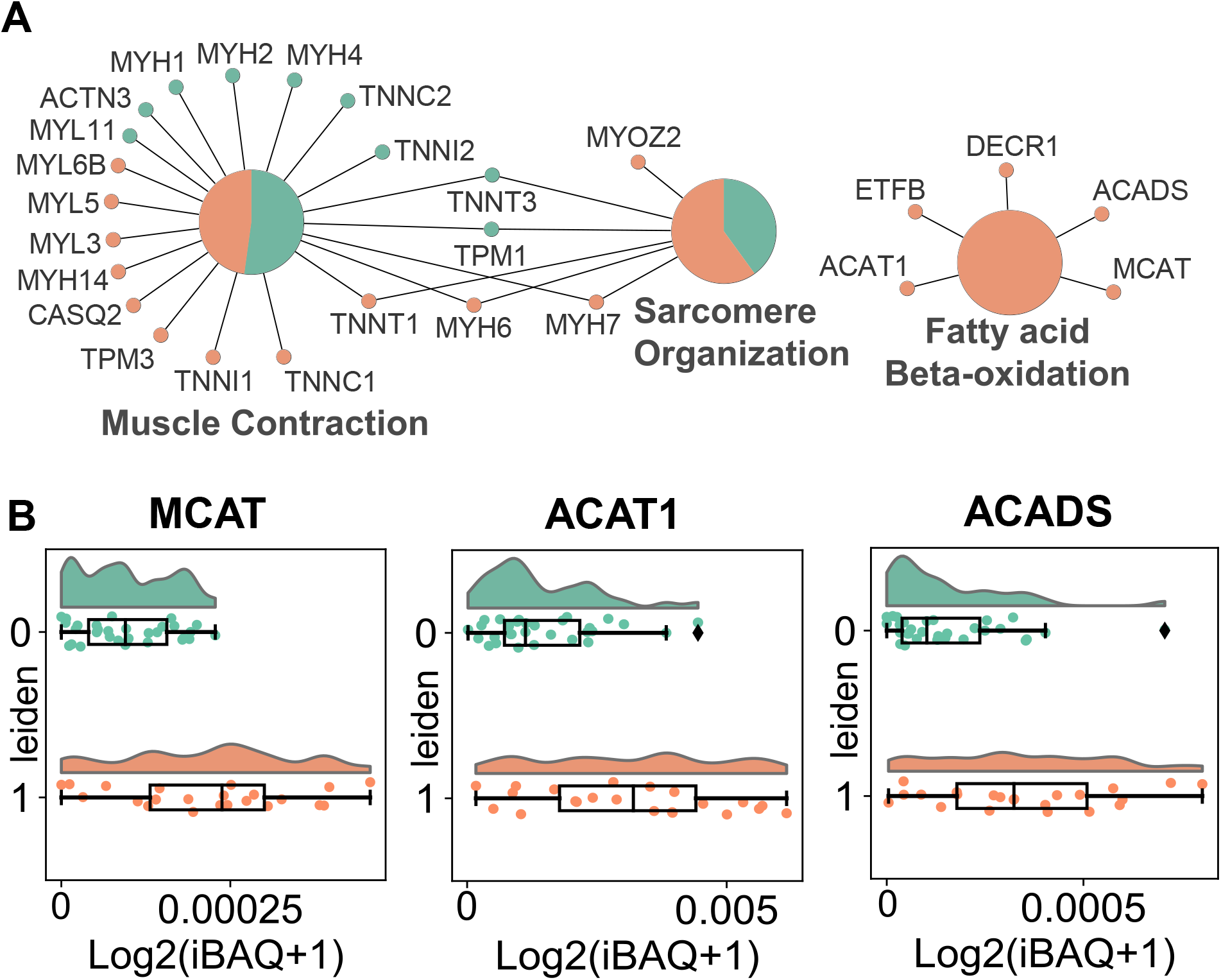
Term enrichment analysis. **(A)** Network of Gene Ontology (GO) Biological Process enrichment of 65 differentially expressed proteins between Leiden clusters 0 and 1 showing two main pathways, muscle contraction and fatty acid beta-oxidation. The large circles indicate enriched GO terms and the small circles show the proteins that are found in those pathways (green: proteins upregulated in Leiden cluster 0; salmon: proteins upregulated in Leiden cluster 1). **(B)** Raincloud plots of fatty acid beta-oxidation proteins that were statistically higher in Leiden cluster 1 compared to 0.

## DISCUSSION

Our high throughput single muscle fiber data acquisition and analysis is able to quantify and distinguish human type 1 and 2A fibers more quickly than previously described methods for skm single fiber proteomics. In ∼1/10th the time, we are able to identify differences in protein quantities that are congruent with established skm physiology. We were also able to adapt a single cell data clustering tool originally designed for single cell transcriptomics data to streamline data interpretation. Furthermore, UMAP and Leiden clustering was able to separate slow and fast fibers, as well as ‘hybrid’ fibers, into distinct groups. When this high throughput mass spectrometry method is paired with a clustering analysis tool, understanding of single skm fiber proteomes on the scale of a several hundred fibers analyzed per individual will be possible, which is a more accurate representation of the whole muscle than the current approach of <20 fibers per person.

Similar to other skm studies^7^, we found most fibers (83%) were composed of a single dominant MYH isoform, namely pure MYH2 (type 2A) and MYH7 (type 1), and only a few fibers had mixed quantities of two MYH subtypes (**Figure 3C**). The mixed subtypes, or ‘hybrid’ fibers, observed were MYH2/7 (1/2A) and MYH2/1 (2A/2X), which are the most commonly observed hybrid fibers in humans when fiber types are determined with immunohistochemistry of muscle cross-sections or SDS-PAGE of isolated single muscle fibers^2^. We found that MYH protein quantities calculated using iBAQ from DIA-NN precursor data with subsequent actin normalization were more reliable in assigning MYH isoforms based on established knowledge of MYH subtype distributions within human skm compared to protein quantities from MaxLFQ. We characterized the differences in type 1 and 2A fiber types based on the quantities of the regulatory and structural proteins (troponins, tropomyosins, MYL proteins, Z-line linking proteins, and a sarcoplasmic reticulum protein) and found a majority of the proteins (**Figure 2 and Figure 5)** to also be congruent with previous findings^3,30^. For example, MYL3, which we found to be significantly higher in Leiden 1 (mostly type 1 fibers), has been previously reported as 10 times more abundant in type 1 versus type 2A fibers^3^. Other MYLs that we found to be differentially expressed in slow fibers, namely MYL6B and MYL5, were also reported to be significantly higher in human type 1 fibers^3^. Additionally, the quantity of ACTN3, an actin binding protein, was previously found to be three times higher in type 2X compared to type 1 and 2A fibers ^3^. Although we did not find 2X fibers in our dataset, we did observe that ACTN3 protein was differentially higher (log_2_ fold change of 2) in Leiden 0 (mostly type 2A) compared to Leiden 1 (mostly type 1). Together, these findings indicate that the depth and sensitivity of our approach was uncompromised by the nearly 10-fold decrease in data acquisition time.

In addition to the expected differences we observed in the regulatory and structural proteins between fiber types, we found five proteins (ACAT1, ETFB, DECR1, ACADS and MCAT) differentially higher in Leiden 1 that play key roles in mitochondrial beta-oxidation, which aerobically breaks down fatty acids to produce ATP. For example, ACADS (short chain specific acyl-CoA dehydrogenase) catalyzes one of the pathway’s first steps and breaks down fatty acids into acetyl-CoA, ETFB (electron transfer flavoprotein subunit beta) receives electrons from several mitochondrial dehydrogenases, and ACAT1 (acetyl-CoA acetyltransferase) catalyzes the last step of this pathway. These findings are consistent with established fiber type differences, as Leiden 1 is composed of primarily type 1 fibers which are well-known to have higher mitochondrial content as they rely more on oxidative phosphorylation rather than glycolysis to generate ATP^1^. An unexpected observation, however, was that we did not find differences in proteins involved in glycolysis between the two Leiden groups, as type 2A fibers are also known to rely more on glycolysis than type 1 fibers. One potential explanation for this observation is that we included fibers from both a young and older woman, and it was recently observed that proteins involved in glycolysis and glycogen metabolism were higher in slow fibers and lower in fast fibers from older compared with younger men^7^. Thus, the inclusion of the older and younger women in our dataset may have masked the expected fiber type differences in glycolytic protein content. Further studies are warranted to investigate if the metabolic protein profile differs between type 1 and 2A fibers from young and older women.

Our data indicates MYH6 (alpha isoform in cardiac muscle), MYH4 (primarily in rodent limb muscles) and MYH14 (a non-muscle myosin) were significantly different between Leiden groups, however the abundance of each of these isoforms was <1% of the total MYH content. This is in line with the current literature, which shows MYH6 and MYH4 to be completely absent or in very low abundance in human skeletal muscle fibers ^1,7,30^. Nevertheless, the mean iBAQ-calculated quantity of MYH6 in our data was 5e-4 in Leiden group 1 and 8e-6 in Leiden group 0, which led to it being significantly different between groups despite the low amount. It is possible that the presence of these myosins is due to the allowed 1% false discovery rate in our dataset. Misassignment of peptide sequences is especially problematic with MYH isoforms, where sequence homology is high; in this case the peptide contributing most of the MYH6 quantity is one amino acid different from MYH7 (a Y in MYH6 versus a F in MYH7). We had variable methionine oxidation off in our search due to the additional time cost in library-free DIA-NN searching; for this peptide, the tyrosine detected could actually be the F amino acid residues with neighboring oxidized M on the peptide from MYH7. This highlights a benefit of using the iBAQ approach for protein quantitation, which is that the reported quantities of proteins that are identified by a single peptide are penalized by the denominator that divides by the number of possible detectable peptides. In this way if these protein IDs are among our 1% false positives, their quantity will be very small because it is unlikely to get multiple false hits to the same protein. Further studies with western blot or immunohistochemistry will reveal if there is in fact a small but significant increase in MYH6 in type 1 relative to type 2A fibers.

An unexpected observation in our study was that actin appeared at a much higher quantity than all other proteins in our dataset (**Figures 1 and 2**). In contrast, a skm proteomics study on mice found myosin to be the most abundant protein, not actin^32^. Upon further investigation, the high abundance of actin in our study appeared to be a consequence of the denominator in the iBAQ protein calculation, which in our implementation divides each protein by the number of potential unique observable peptides as a normalization of the summed peptide signal numerator. Specifically, actin is substantially shorter than myosins (377 amino acids in ACTS versus 1,935 amino acids in MYH7), and thus, the denominator used in the iBAQ calculation for ACTS was 7, while the denominator used for MYH7 was 120. This resulted in a difference in the ratio of unique observable peptides to amino acids to be 7/377 = 0.02 for ACTS compared with 120/1,935 = 0.06 for MYH7. Therefore, the normalization ratio for ACTS was skewed relative to the protein’s length due to many non-unique peptides in ACTS. This does not adversely affect the results of this study because we only compared ratios between proteins with similar properties (e.g., myosins and tropomyosins). It is also generally well accepted that comparing ratios between different proteins from label free quantification proteomics data provides a ballpark estimate at best due to the differences in peptide ionization efficiency.

## CONCLUSION

We report a complete workflow for high throughput single muscle fiber proteomics including sample preparation, data collection, and bioinformatic summary analysis. We also demonstrate that the iBAQ quantification algorithm for protein quantification better reflects the myosin heavy percentages based on known muscle physiology than those derived from MaxLFQ. The proteome data cleanly clusters itself into two groups that are mainly driven by MYH types. Comparing the two groups from clustering showed protein differences between the groups that are consistent with known muscle physiology. In the future our workflow will enable large scale studies of human muscle fiber heterogeneity.

## SUPPORTING INFORMATION

**Table S1**. iBAQ-calculated protein quantities normalized by quantity of ACTS per fiber for all 744 proteins and 53 fibers.

**Table S2**. Fraction of each MYH isoform to total MYH per fiber calculated using protein quantities from iBAQ.

**Table S3**. Fraction of each MYH isoform to total MYH per fiber calculated using MaxLFQ.

**Table S4**. Means, log2(fold change) and -log10(B-H adjusted p-value) for all 744 proteins between Leiden clusters 0 and 1. Positive log2(fold change) means protein quantity was higher in Leiden cluster 0 compared to 1, negative log2(fold change) means protein quantity was higher in Leiden cluster 1 compared to 0.

**Figure S1**. Ten most abundant proteins in slow and fast fibers when quantities are derived from MaxLFQ.

**Figure S2**. iBAQ-calculated MYH fractions prior to removal of MYH4 peptide with overlap with MYH2 and description of peptide that was removed.

**Figure S3**. Visualization of TPM1 and TPM3 per fiber and correlations with fast and slow fibers.

**Figure S4**. Volcano plot highlighting 65 differentially expressed proteins between leiden clusters and 1.

**Figure S5**. UMAP colored by old and young fiber types.

## ACKNOWLEDGEMENTS

This study was supported by NIH grants R21AG074234 and R35GM142502 to JGM and R01AG048262 to CWS. We thank H. Adam Steinberg for graphic design assistance.

## AUTHOR CONTRIBUTIONS

Conceptualization, A.M., J.G.M., C.W.S., and A.Hu.; Methodology, A.M., J.G.M., A.Hu., S.K., and Y.J.; Software, A.M., J.G.M., and A.Hu.; Formal Analysis, A.M. and J.G.M.; Investigation, S.K., Y.J., A.Ha., M.M., S.Y., L.E.T., and C.S.Z.; Resources, J.G.M., C.W.S., J.V.E., and S.J.P.; Data Curation; A.M. and J.G.M.; Writing - Original Draft, A.M. and J.G.M.; Writing - Review & Editing; A.M., Y.J., S.K., L.E.T., C.S.Z., A.Ha., M.M., S.Y., A.Hu., S.J.P., J.V.E., C.W.S., and J.G.M.; Visualization, A.M.; Supervision, J.G.M., C.W.S, J.V.E., and S.J.P.; Projection Administration, J.G.M. and C.W.S.; Funding Acquisition, J.G.M. and C.W.S.

