## Supplementary Figures for "Complete Workflow for High Throughput Human Single Skeletal Muscle Fiber Proteomics"

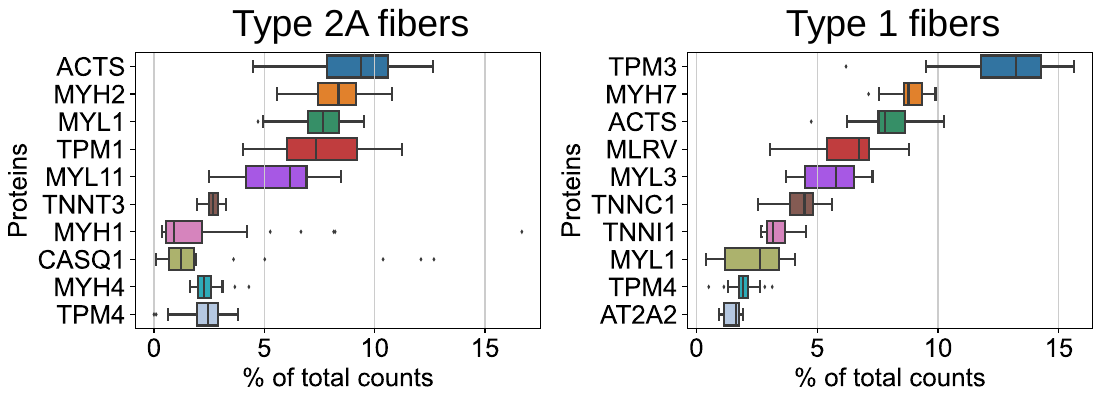

**Figure S1 Ten most abundant proteins when quantities are computed by MaxLFQ.** Top ten proteins with highest percent of total counts in type 1 (left) and type 2A (right) fibers using MaxLFQ quantification. MLRV is interchangeable with MYL2. MYL11 is interchangeable with MYLPF and MLC2-fast.

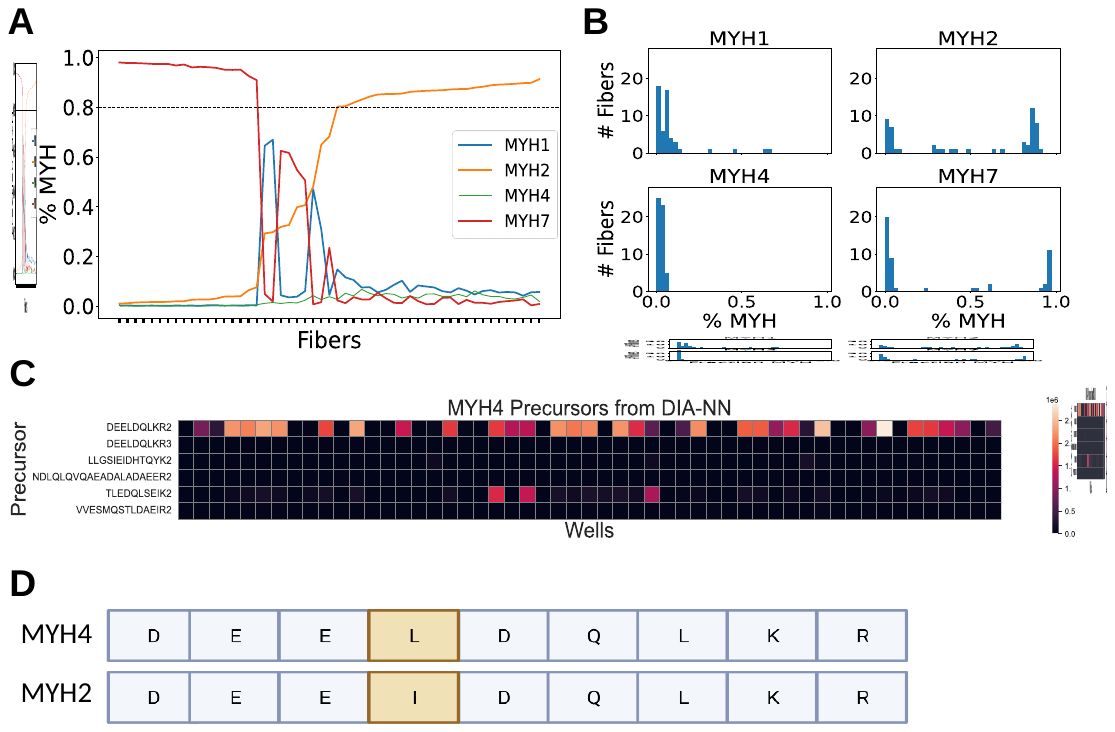

**Figure S2 MYH4 peptide overlap with MYH2. (A)** Fraction of MYH subtypes (sorted by MYH2 from low to high) using protein quantities from iBAQ prior to removing MYH4 peptide with overlap with MYH2. **(B)** Histogram showing MYH fractions per fiber using protein quantities from iBAQ prior to removing MYH4 peptide with overlap with MYH2. **(C)** Precursors mapping to MYH4 with one peptide driving the abundance of MYH4. Peptide charge is given at the end of each sequence. **(D)** BLAST results for DEELDQKR^2+^ peptide indicate identical sequence to MYH2 except for substitution of I for L.

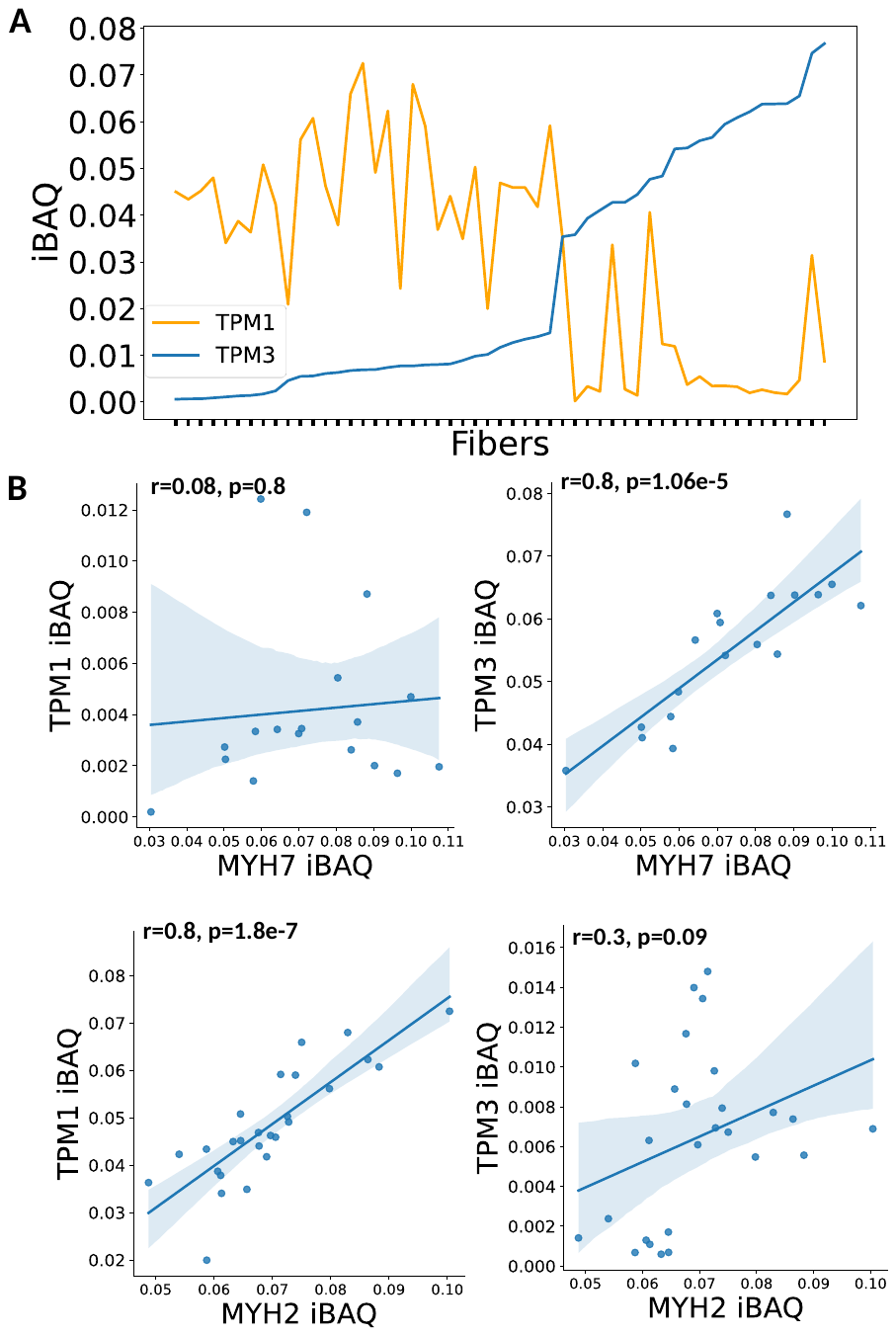

**Figure S3 TPM1 and TPM3 quantities per fiber and correlation with fast and slow fibers. (A)** iBAQ-calculated quantity of TPM1 and TPM3 by fiber sorted by TPM3 values from low to high. **(B)** Linear regression plots visualizing relationships between MYH2, MYH7 and TPM1, TPM3 and corresponding Pearson correlation coefficients (r) for each plot.

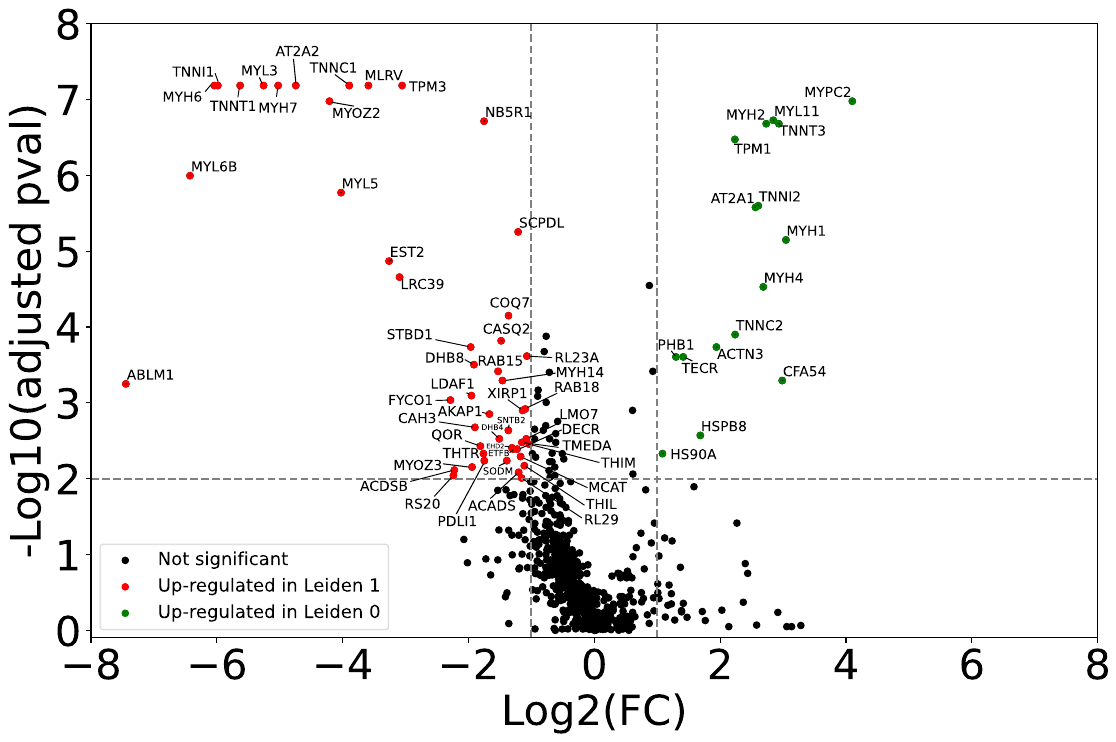

**Figure S4** **Volcano plot depicting all proteins.** Proteins above log_10_(adjusted p-value) 2 (equivalent to B-H adjusted p-value below 0.01) are proteins that are statistically different between Leiden cluster 0 and 1. Proteins to the right of log_2_(fold change of Leiden 0/Leiden 1) 1 and left of -1 represent fold changes of Leiden 0/Leiden 1 greater than or less than 2, respectively. Sixty-five proteins were defined as significantly different according to the adjusted p-value cutoff of < 0.01 and absolute value of log2(fold change) >1 (**Supplementary Table 4**).

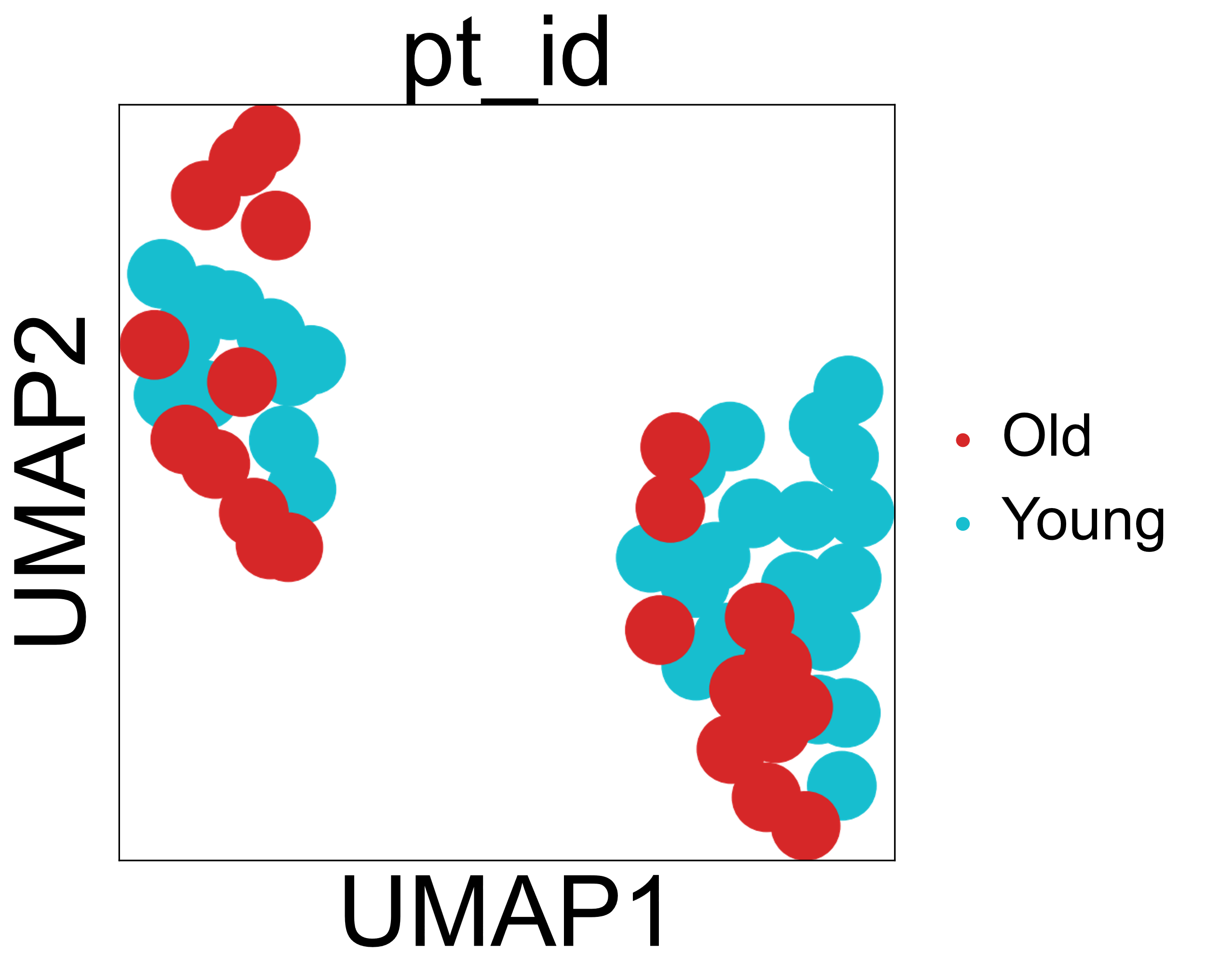

**Figure S5 UMAP distribution of old and young fibers.** UMAP in the same layout as Figure 3 colored by young and old fibers.
